# Cross-kingdom signalling regulates spore germination in the moss *Physcomitrella patens*

**DOI:** 10.1101/839001

**Authors:** Eleanor F. Vesty, Amy L. Whitbread, Sarah Needs, Wesal Tanko, Kirsty Jones, Nigel Halliday, Fatemeh Ghaderiardakani, Xiaoguang Liu, Miguel Cámara, Juliet C. Coates

## Abstract

Plants live in close association with microorganisms that can have beneficial or detrimental effects. The activity of bacteria in association with flowering plants has been extensively analysed. Bacteria use quorum-sensing as a way of monitoring their population density and interacting with their environment. A key group of quorum sensing molecules in Gram-negative bacteria are the *N*-acylhomoserine lactones (AHLs), which are known to affect the growth and development of both flowering plants, including crops, and marine algae. Thus, AHLs have potentially important roles in agriculture and aquaculture. Nothing is known about the effects of AHLs on the earliest-diverging land plants, thus the evolution of AHL-mediated bacterial-plant- and algal interactions is unknown. In this paper, we show that AHLs can affect spore germination in a representative of the earliest plants on land, the Bryophyte moss *Physcomitrella patens*. Furthermore, we demonstrate that sporophytes of wild isolates of *Physcomitrella patens* are associated with AHL-producing bacteria.

## Introduction

Plants do not exist in isolation in the environment, but interact with a wide array of organisms from all kingdoms including bacteria, fungi, animals and other plants. These interactions can have profound effects on plant fitness, growth and development. In addition to pathogenicity or parasitism, interactions between plants and other organisms can be beneficial. Examples include interactions between fungi and plants in the form of mycorrhizae and interactions between plants and bacteria ^1–3^.

Interactions between plants and microorganisms have become more elaborate during plant evolution. Mycorrhizal interactions are beneficial in liverworts, one of the earliest diverging groups of land plants, where the association between liverworts and fungi boosts plant photosynthesis, growth, fitness and nitrogen/phosphorus uptake ^4,5^. In later-diverging plants including flowering plants, interactions with microorganisms have increased in complexity. For example, in legumes (Fabaceae, including beans and pulses) plant root cells are surrounded by a group of proteobacteria (Rhizobia) and form a symbiotic root nodule ^6^ while actinobacteria from the *Frankia* genus can form nodules with a wide range of plant families ^7,8^. The bacteria gain carbon from the plant, while the plant gains nitrogen from the bacteria. Not all plant-bacterial interactions are so highly specialised: many bacteria in the rhizosphere contribute to plant productivity and gain from plants in return ^9^.

It has become clear that flowering plants can respond to bacterial signalling molecules that alter plant growth and development, representing inter-kingdom interaction ^10,11^. The perception of bacteria by plants is of significant importance in terms of monitoring their surroundings and thus being able to respond accordingly to enhance their chances of survival ^12^.

A key way in which bacteria communicate with one another is via diffusible quorum sensing (QS) molecules that are used to monitor and respond to population density within a colony or biofilm ^13–19^. One well-characterised subset of QS molecules that affect plants behaviour are the *N*-acylhomoserine lactones (AHLs) ^11,12,20–24^. AHLs are produced by Gram-negative bacteria ^25^ and are key for the control of multiple gene expression in a coordinated manner within a population ^25–27^.

AHLs vary in their structure in nature with a wide range of acyl chain lengths, from four to eighteen carbons, and level of saturation. Furthermore, at the third carbon position (C3), different substitutions can also occur whereby the molecule can either be unsubstituted, contain a ketone group (oxo) or a hydroxyl group (OH). These structural differences contribute to their specific impact on gene expression ^28,29^.

A wide range of Gram-negative plant-associated bacteria produce AHLs and some non-producers are still able to sense and respond to the presence of these molecules using *luxR*-solo AHL receptor proteins ^30,31^. The ability of plants to detect and respond to the presence of AHLs may be a result of their coevolution with AHL-producing bacteria. Plant perception of AHLs may provide an evolutionary advantage over their associated microbial community, especially if the bacteria are pathogenic, enabling the plants to detect increasing bacterial populations and alter the QS outcome ^32^. Plant responses to AHLs are dependent on the structure and concentration of the AHL encountered and can be positive or negative in terms of growth ^33,34^. Plant-bacterial interaction often occurs in the rhizosphere where roots in the soil come into contact with AHLs in varying concentrations due to bacterial growth ^21,35,36^. Flowering plants respond to these bacterial compounds and even absorb them from the surrounding environment ^37^. The role of QS in legume nodule formation seems to vary depending on the combination of plant and bacterium under investigation (reviewed in ^34^).

Certain plant species produce AHL mimics that induce a premature quorum-sensing response in bacteria that serves to protect the plant from pathogens, or aid establishment of symbiotic relationships ^38–42^. Conversely, plants can produce anti-QS molecules and use “quorum quenching” to interfere with bacterial QS signalling mechanisms preventing transcription of specific gene sets, thus averting the synthesis of virulence factors by pathogenic bacteria ^43,44^. The exact mechanisms by which plants perceive AHLs is currently unknown, but these molecules can affect the activity of endogenous plant signalling, such as calcium signalling ^45,46^, G-protein signalling ^47,48^, stress signalling and metabolism ^23,49^ and hormone signalling ^22^ causing downstream effects on plant growth.

In plants, AHL perception induces a “primed” state, regulating plant immunity ^12^. Tomato plants (*Solanum lycopersicum*) became resistant to *Alternaria alternata*, a pathogenic fungus, following co-culture with wild-type *Serratia liquefacians* bacteria, whereas no induced resistance occurred following the growth of a corresponding *Serratia* AHL-deficient mutant ^50^. In the model plant *Arabidopsis thaliana*, resistance against both bacterial and fungal pathogens was observed following the exogenous application of synthetic AHLs ^12^. However, it is not just plant immunity that is affected by QS signal molecules: AHLs also affect plant growth, development and physiology. Addition of AHL to plants modifies protein profiles by inducing changes in gene expression ^45,51^, altering the formation of roots ^11,20–23^, including the promotion of adventitious root growth ^24^. Plant-growth-promoting bacteria employ QS systems which aid with the colonisation of the rhizosphere, providing benefit for both bacteria and host plant ^52,53^. The ecology of rhizosphere bacterial populations on *S. lycopersicum* is modulated by AHLs produced by bacteria a significant distance away, due to the movement of the compounds through the rhizosphere via diffusion ^14,54^. Furthermore, there are potential beneficial effects of the by-products of AHL degradation. Exogenous application of homoserine lactones, and homoserine, the degradation products of AHLs, to bean roots increased the stomatal conductance of the plants, which in turn led to enhanced mineral nutrient availability, benefiting both the host plant and rhizosphere-associated bacteria ^55^.

Plant-produced compounds, such as strigolactones and alkamides, share structural similarity to the bacteria-generated AHL molecules. Consequently, it is not surprising that QS molecules impose effects on the growth and development of plants, as both alkamides and strigolactones are known to induce a number of morphological responses, including changes in root architecture ^21^. An intact homoserine lactone ring structure is not always required for plants to detect AHLs ^33^.

All plants on land arose from aquatic ancestors: the appearance of plants on land was a key evolutionary transition. Relatively little is known about the effects bacteria have on development in ancient plant lineages ^56^. The earliest land plants were small and in close contact with their substrate (and associated microorganisms) over the whole of their anatomy rather than just via their roots. The earliest-diverging lineage of land plants, the spore-bearing mosses, liverworts and hornworts (Bryophytes) play a key role in ecology as carbon sinks in peat bogs and permafrosts ^57^ and have been used by humans for their absorptive and medicinal properties for thousands of years ^58–60^.

The microbiome of *Sphagnum* moss harbours diverse bacteria and is substantially different from that of flowering plants with the potential to enable plant- and ecosystem adaptation to climate change ^56,61–63^. The microbiomes of a co-occurring epiphytic moss (*Pterygynandrum filiforme*) and its flowering plant host (*Acer pseudoplatanis*) show distinct characteristics ^64^. Moreover, different moss species from different habitats possess distinct microbiomes with some overlap in properties and function ^65,66^.

Whether bacterial signalling molecules can directly affect developmental processes in non-flowering plants is largely unknown. A symbiotic bacterium (*Methylobacterium*) from the moss *Funaria hygrometrica* exerted a cytokinin-like effect on moss development, enabling formation of buds, and promoted filament growth via cell division ^67,68^. Evidence from marine seaweeds (macroalgae), which share a common ancestor with land plants, demonstrates that AHLs from algal-associated bacteria can affect algal growth, development and cell behaviour ^69–72^. Motile, reproductive spores of the green seaweed *Ulva* sense and are attracted to AHLs produced by bacterial biofilms, which influence spore settlement and swimming rate ^71–73^ and cause activation of calcium signaling in the spores ^72^. AHLs from *Shewanella* and *Sulfitobacter* inhibit early development of the green seaweed *Ulva* from spores and synthetic *N*-dodecanoyl-L-homoserine lactone (C12-HSL) inhibits early Ulva development at concentrations above 5µM ^69^. AHLs (*N-*butanoyl-L-homoserine lactone (C4-HSL) and *N-*hexanoyl-L-homoserine lactone (C6-HSL)) from *Shewanella* promote reproductive carpospore release in the red seaweed *Gracilaria dura* at micromolar concentrations ^70^.

We therefore hypothesised that AHLs might affect development in early-diverging land plants. In this paper, we show that synthetic AHLs can promote spore germination in the model moss species *Physcomitrella patens* ^74,75^ in a lab-based assay. Moreover, sporophytes from wild isolates of *Physcomitrella patens* are associated with AHL-producing bacteria, suggesting these bacteria may influence spore germination in the environment through the production of AHL signal molecules.

## Results

### AHLs promote *Physcomitrella* spore germination at sub-micromolar concentrations but inhibit spore germination at concentrations above 1µM

Previous studies have shown that AHLs at concentrations of 1-10µM can promote root growth in the model flowering plant *Arabidopsis* (Jin et al., 2012; Liu et al., 2012; vonRad et al., 2008; Zhao et al., 2015). In algae, AHLs at 2-10µM can promote spore release ^70^ or reduce the progress of growth and development from spores ^69^. We tested the effect of a range of AHLs with different carbon *N*-acyl chains lengths at 0.1µM and 1µM concentrations on the spore germination of the model moss *Physcomitrella patens*. All AHLs (C4-HSL to C12-HSL) induce a significantly faster spore germination rate compared to a solvent-only control (Figure 1). C4-HSL and C6-HSL show a similar promotion of germination at both 0.1µM and 1µM concentrations (Figure 1A, B). C8-HSL appears to have slightly more germination-promoting activity than the shorter chain AHLs and is more potent at 1µM than 0.1µM (Figure 1C). C10-HSL and C12-HSL are the most potent germination-promoting AHLs, being more effective at 0.1µM than at 1µM concentration (Figure 1D, E).

**Figure 1.**
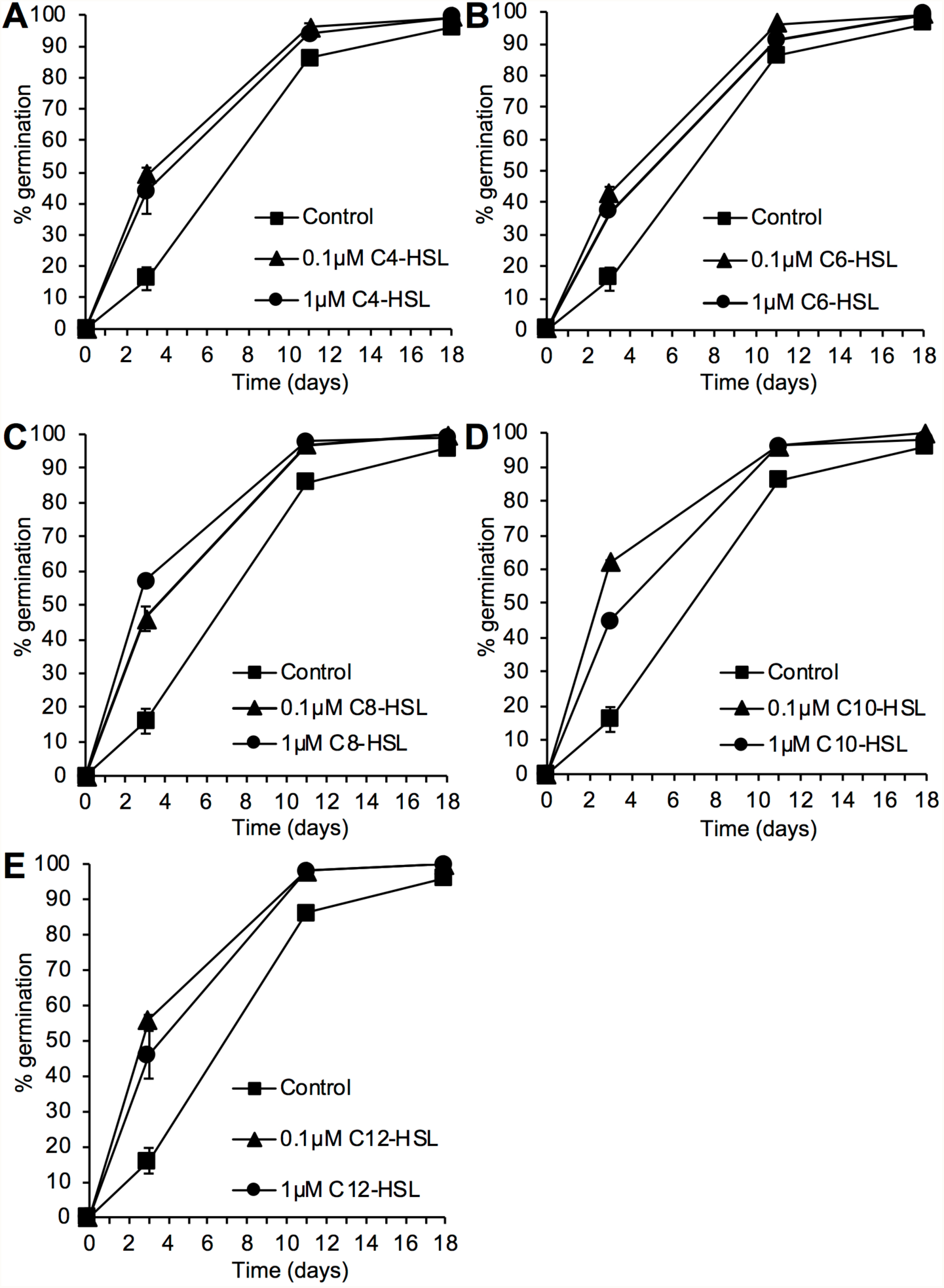
*N*-acyl HSLs can promote *Physcomitrella* spore germination. *P. patens* spores were germinated on media containing 0, 0.1 and 1µM *N*-acyl HSLs of varying chain lengths (C4-C12). The number of spores germinated were counted as a percentage of total spores on the plate. Both concentrations of *N*-acyl HSLs resulted in a faster rate of germination compared to control spores. A) C4-HSL promotes germination of *P. patens* spores. Z-tests indicated significant differences in germination between treated and untreated spores on days 3 and 11 (P >|t| 0.0002). B) C6-HSL promotes germination of *P. patens* spores. Z tests indicated significant differences in germination between treated and untreated spores on days 3 and 11 (P >|t| 0.0002). Treatment with the lower concentration of 0.1µM was more effective in promoting germination on days 3 and 11 when compared to 1µM. C) C8-HSL promotes germination of *P. patens* spores. Z tests indicated significant differences in germination % between treated and untreated spores on days 3, 7 and 11 (P >|t| 0.0002). D) C10-HSL promotes *P. patens* spore germination. Z tests indicated significant differences in germination % between treated and untreated spores on days 3, 7 and 11 (P >|t| 0.0002). Treatment with the lower concentration of 0.1µM was significantly more effective in promoting germination on days 3 and 7 when compared to 1µM. E) C12-HSL promotes *P. patens* spore germination. Z tests indicated significant differences in germination % between treated and untreated spores on days 3, 7 and 11 (P >|t| 0.0002). Treatment with the lower concentration of 0.1µM was significantly more effective in promoting germination on days 3 and 7 when compared to 1µM. In all experiments, final germination efficiency was not affected with all treatments achieving a final germination of over 95%. Representative of more than 5 biological repeats. Error bars represent ± SEM.

In *Arabidopsis*, 50-100µM concentrations of AHLs inhibit root growth ^21,45,48^ while in the seaweed *Ulva* spore germination and early development is reduced with just 5µM AHLs ^69^. We tested the effects of a range of AHLs at 5µM on *Physcomitrella* spore germination and found that AHLs could inhibit spore germination (Figure 2A). The effect appeared strongest with C10-HSL, which also inhibited germination at 10µM, in a dose-dependent manner (Figure 2B). Taken together, these data show that AHLs, particularly those with longer chain length, accelerate spore germination when at low (≤1µM) concentrations and inhibit spore germination at higher (5-10µM) concentrations.

**Figure 2.**
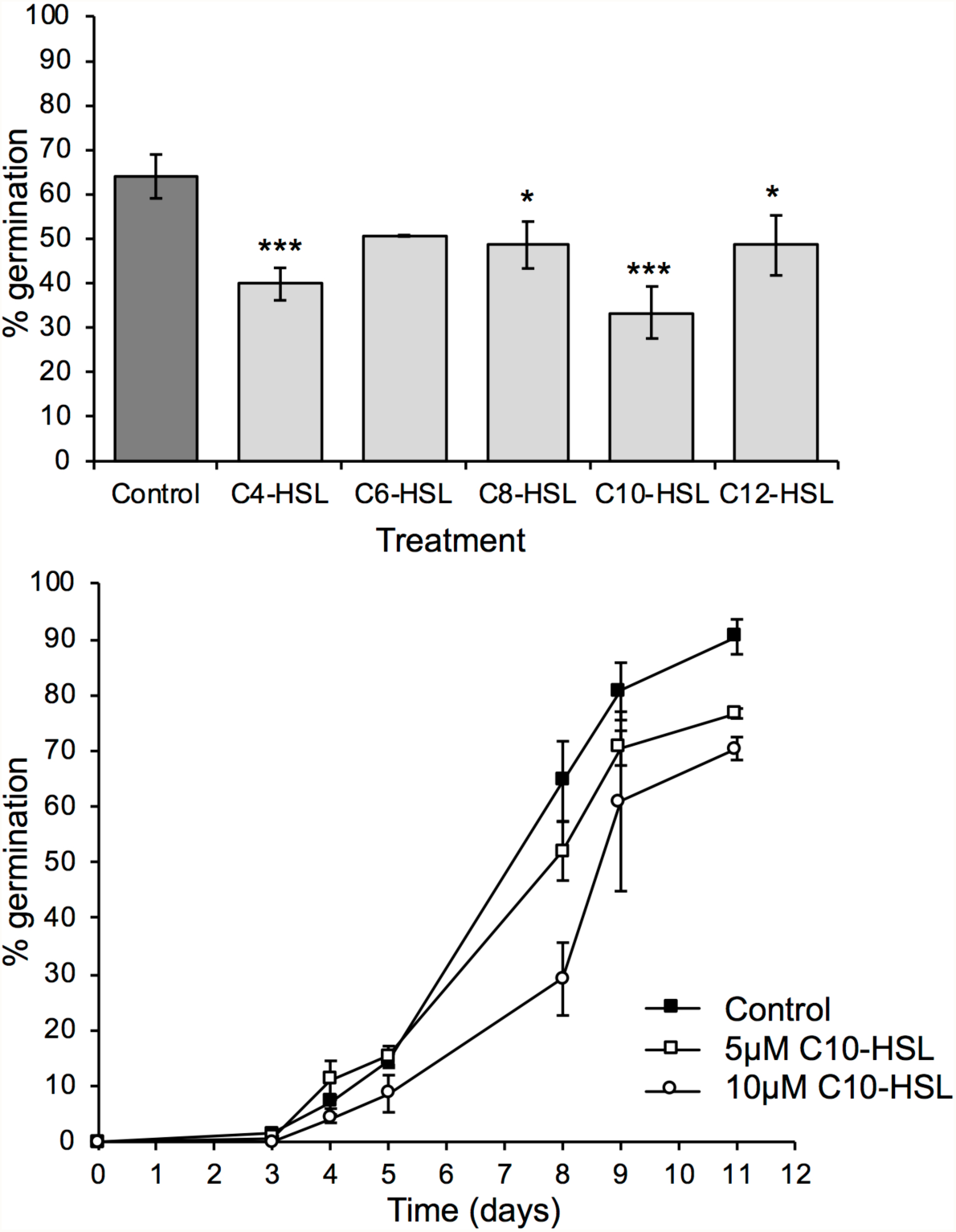
*N*-acyl HSLs inhibit *Physcomitrella* spore germination at concentrations above 1µM. A) C4-C12 *N*-acyl HSLs were tested on *P. patens* spores at a concentration of 5µM (light grey bars) compared to a solvent-matched control (dark grey bar). A snapshot of data at day 4 is shown: all chain lengths reduce germination. Significant differences between control and treatment are seen with a Z-test for C4-HSL (p=0.0007), C8-HSL (p=0.0324), C10-HSL (p<0.0002) and C12-HSL(p=0.0324) but not C6-HSL (p=0.0629). * p<0.05, *** p<0.001. Error bars represent ±SEM. n>700 spores for each data point. Representative of at least 3 biological repeats. B) C10-HSL inhibits *P. patens* spore germination in a dose-dependent manner. C10-HSL was tested at 5µM and 10µM concentration against a solvent control. Significant differences are seen with a Z-test between control and both 5µM and 10µM C10-HSL on day 8, 9 and 11 (p<0.0002); 5µM and 10µM C10-HSL are also significantly different from each other on day 8 (p<0.0002), day 9 (p<0.0002) and day 11 (p=0.0056). Error bars represent ±SEM. n>500 spores for each data point. Representative of 3 biological repeats.

### Chain length and side group substitution affect the activity of AHLs against *Physcomitrella* spore germination

To investigate whether changing the side group of the AHL had an effect on biological activity, we assayed the spore-germination-promoting activity of C4-C12 AHLs, namely the *N*-acyl version (as before) and also the 3-oxo (3-O) and 3-hydroxy (3-OH) substituted forms. Our “snapshot” data (Figure 3A) indicated potential differences in potency between the different side chains, particularly for AHLs with longer carbon chain. To investigate these differences further, we assayed spore germination in the presence of 3-OH and 3-O substitutions of the C10 and C12 HSLs, which consistently through this study showed some of highest activity, over a range of concentrations from 2nM to 1µM (summarised in Figure 3B; data in Supplemental Figure 1). The *N*-acyl variants of C10- and C12-HSL showed the greatest spore germination-promoting activity at 2-10nM (Figure 3B; Supplemental Figure 1A and 1D). 3-OH-C10-HSL variant showed greatest spore germination promotion at 10nM whereas the 3-OH-C12-HSL showed similar spore germination-promotion from 10nM-1µM, slightly higher at 1µM (Figure 3B; Supplemental Figure 1B and 1E). The 3-O variants of C10- and C12-HSL showed greatest spore germination-promoting activity at 0.1µM concentration, indicating somewhat reduced potency compared to the other two types of AHLs (Figure 3B; Supplemental Figure 1C and 1F). These data demonstrate that both chain length and side group substitution can affect the biological activity of exogeneously-applied synthetic HSLs on *Physcomitrella* spore germination.

**Figure 3.**
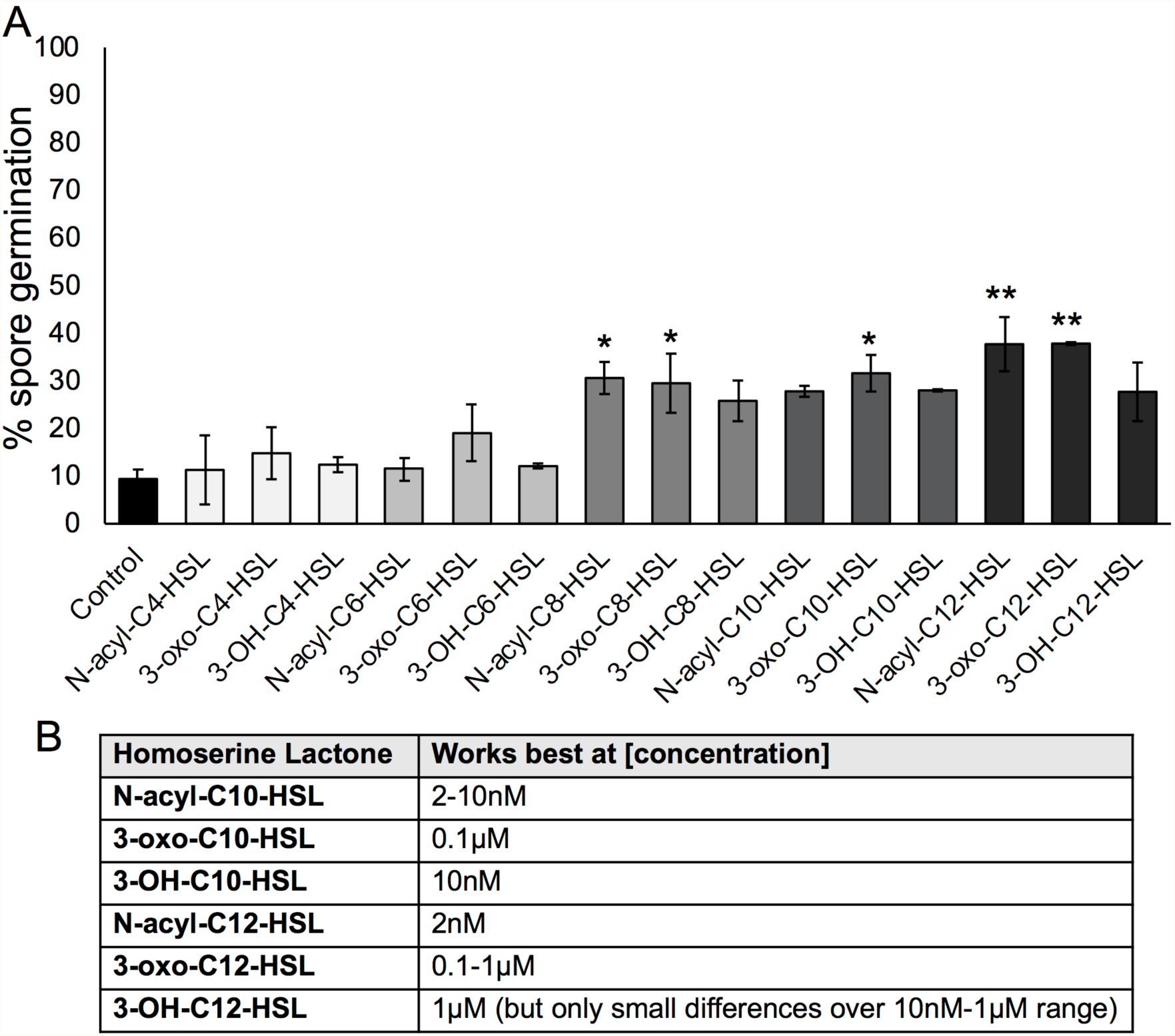
Side chain substitutions affect AHL activity during *Physcomitrella* spore germination. A) 0.1µM of each HSL (*N*-acyl, 3-O or 3-OH) for C4-C12 chain length was tested against solvent control for effects on spore germination. A “snapshot” of germination on day 3 is shown. Asterisks represent significant (*p<0.05; **p<0.01) differences between a treatment and solvent control using a Kruskal-Wallis test and a Dunn’s post-hoc test. Generally, longer chain AHLs stimulate germination more, and AHLs without or with 3-O substitutions appear more potent than those with 3-OH substitutions at this concentration. B) Summary of the optimal concentrations of AHLs for promoting *Physcomitrella* spore germination: full data is shown in Supplemental Figure 1.

### Sporophytes from wild isolates of *Physcomitrella* are associated with multiple species of bacteria, some of which produce AHLs

To determine whether the observed effects of synthetic AHLs on *Physcomitrella patens* spore germination in a lab-based assay might have relevance to wild populations found in the environment, sporulating *Physcomitrella* plants were firstly collected from 3 different locations in the UK with a view to determine the bacterial populations associated with them and the ability of these bacteria to produce AHLs. A total of 12 plants were selected from each sampled location and the sporophyte from each plant was isolated to enable isolation of its associated bacteria (a consortium from each sporophyte). Bacterial consortia were obtained from 30 out of 36 sporophytes. Each individual sporophyte’s (assumed mixed) bacterial populations were taken through multiple rounds of streaking to isolate individual strains associated with *Physcomitrella*. To identify each strain, a fragment of the 16S rRNA gene was amplified from genomic DNA and sequenced. Our sampling of bacteria associated with *Physcomitrella* sporophytes identified largely Proteobacteria from the class Gamma-proteobacteria. Bacteria of the genus *Pseudomonas* were found at all 3 sites (at least 5 different species), as was *Stenotrophomonas* (2 species). Serratia (2 species) were isolated from 2 sites and *Acinetobacter, Aeromonas* and *Rahnella* were each recovered from a single site. The gram-positive bacteria *Microbacterium* (Actinobacteria) and *Bacillus* (Firmicutes) were each found at a single site (Table 1).

**Table 1.**
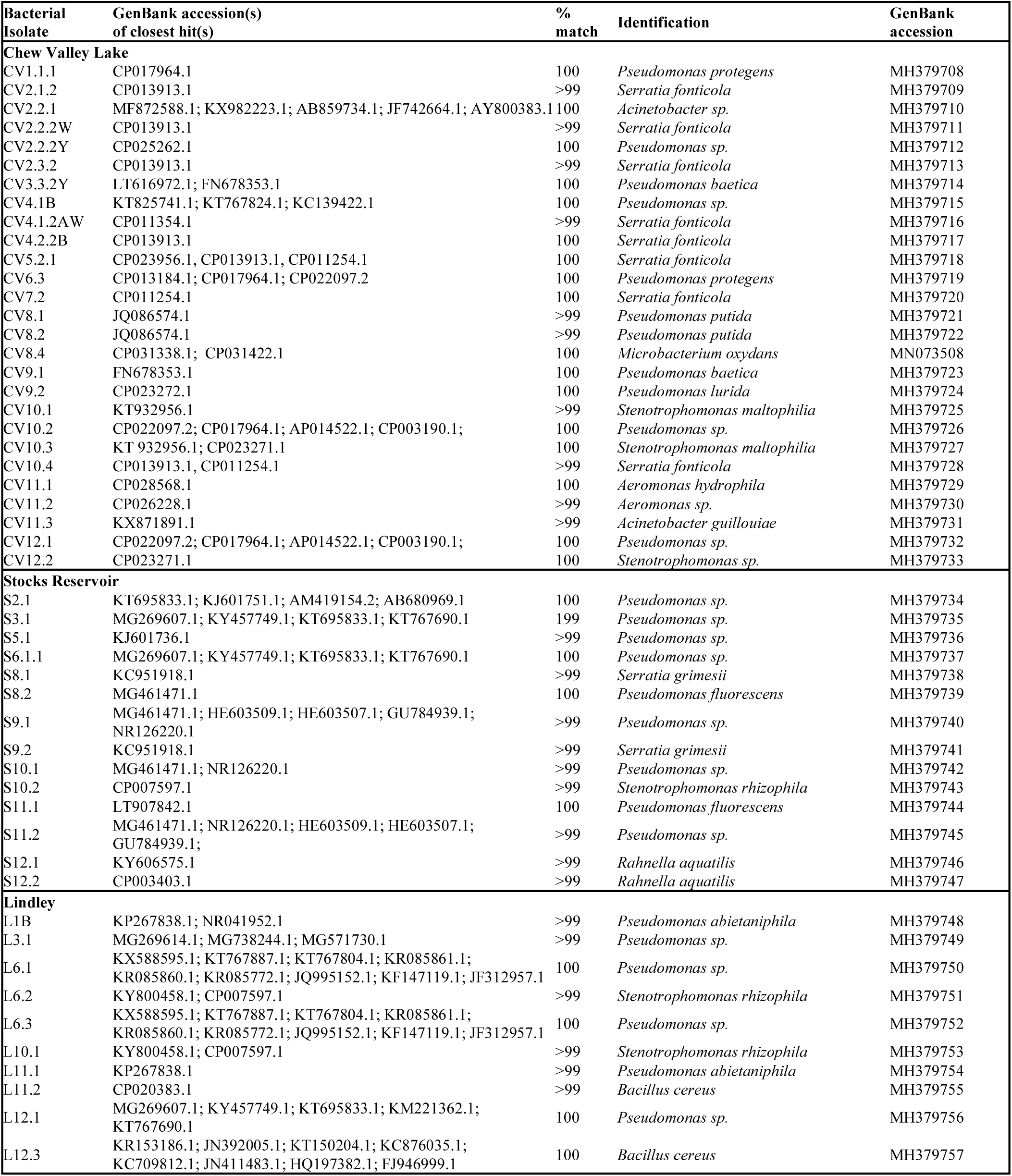
Identification of bacterial isolates from Chew Valley, Stocks Reservoir and Lindley using 16S rDNA sequencing. Closest hits by BLAST, percentage identity and identification are shown for each isolate, along with the newly assigned GenBank accession number for each isolate.

For an initial survey of whether the isolated bacterial consortia from each sporophyte could produce AHLs, consortia were analysed by a mass-spectrometry (LC-MS/MS) approach, which demonstrated that consortia from all three locations could produce AHLs although these were detected to only a limited extent from the Lindley site (Table 2).

**Table 2.** Frequency of AHL detection in bacterial consortia. The number of times a particular AHL was detected in the consortium from a single isolated sporophyte is recorded.

|  | Chew Valley (11 sporophytes' consortia) | Stocks (11 sporophytes' consortia) | Lindley (8 sporophytes' consortia) |
| --- | --- | --- | --- |
| C4-HSL | 2 | 0 | 0 |
| C6-HSL | 0 | 7 | 0 |
| C8-HSL | 8 | 1 | 0 |
| C10-HSL | 0 | 0 | 0 |
| C12-HSL | 0 | 0 | 0 |
| C14-HSL | 0 | 0 | 0 |
| 3-O-C4-HSL | 0 | 0 | 0 |
| 3-O-C6-HSL | 0 | 7 | 0 |
| 3-O-C8-HSL | 1 | 1 | 0 |
| 3-O-C10-HSL | 0 | 0 | 0 |
| 3-O-C12-HSL | 0 | 0 | 0 |
| 3-O-C14-HSL | 0 | 1 | 0 |
| 3-OH-C4-HSL | 0 | 0 | 0 |
| 3-OH-C6-HSL | 0 | 2 | 1 |
| 3-OH-C8-HSL | 7 | 1 | 1 |
| 3-OH-C10-HSL | 8 | 1 | 1 |
| 3-OH-C12-HSL | 0 | 0 | 0 |
| 3-OH-C14-HSL | 0 | 0 | 0 |

To determine whether the individual bacteria isolated from wild *Physcomitrella* could produce detectable AHLs, cultures of the Gram-negative bacterial isolates were subjected to AHL analysis by LC-MS/MS. Just under half of the bacterial isolates from each of Chew Valley and Stocks reservoir produced detectable AHLs, while no AHLs were detected from the bacteria from Lindley. Representatives of *N*-acyl, 3-O and 3-OH from C4 to C10 chain length were detected (Figure 4). Overall, the most frequently detected AHLs were C6-HSL and 3-O-C8-HSL. The most frequently detected AHL in bacteria from Chew Valley was 3-OH-C10-HSL, whereas that from Stocks was 3-O-C6-HSL.

**Figure 4.**
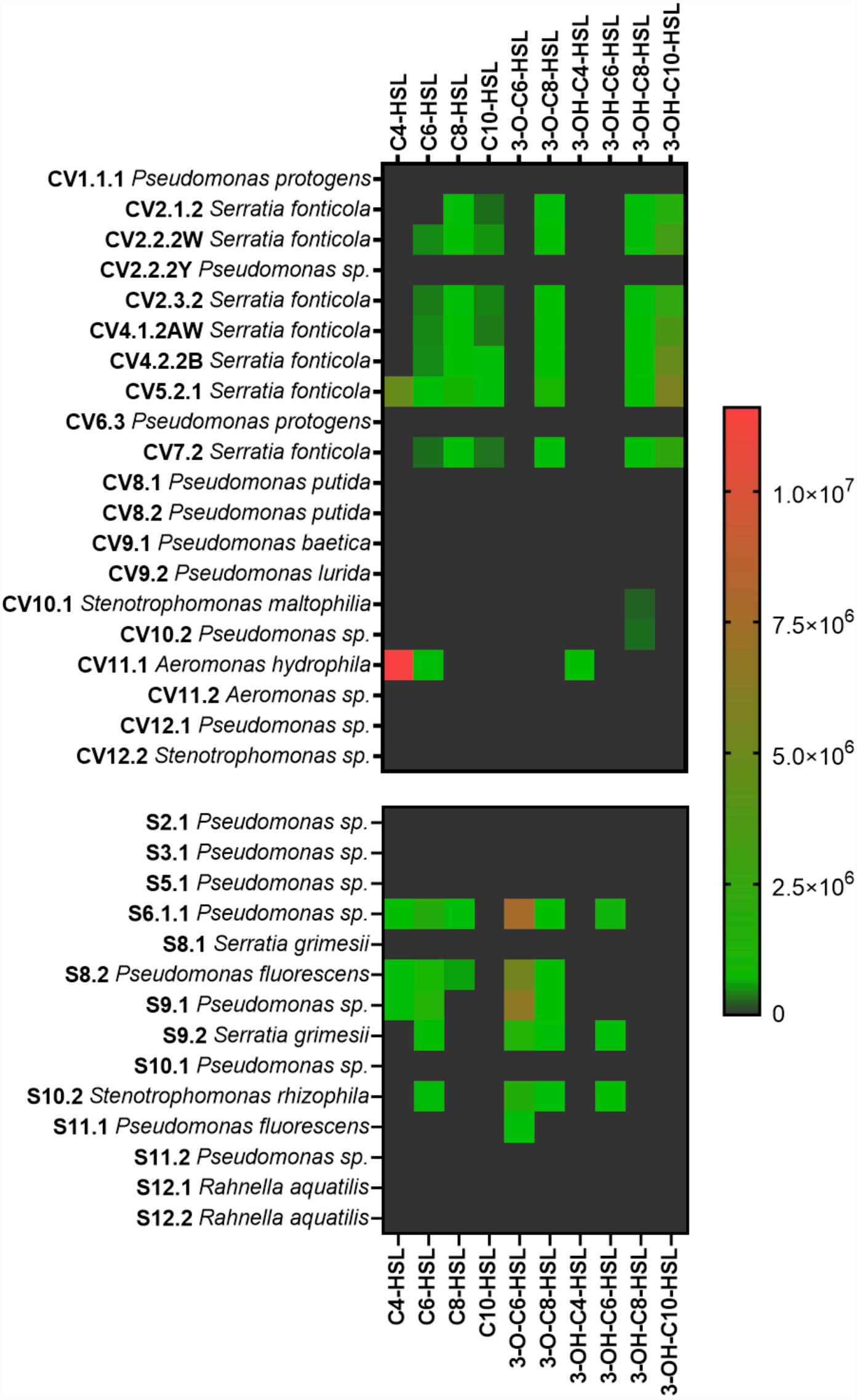
AHLs detected in bacterial isolates from Chew Valley and Stocks Reservoir. No AHLs were detected from individual Lindley isolates for which we obtained high quality sequence. Numerical values on the legend scale are of peak area for detected analytes. A positive detection of an AHL was considered as a chromatographic peak that has a signal to noise ratio of at least 5, displaying a peak retention time that matched that of authentic AHL synthetic standards.

Taken together, these data show that some of the bacteria associated with *Physcomitrella* sporophytes from different geographical locations can produce a range of AHLs.

## Discussion

Our experiments show for the first time that synthetic AHLs can affect the spore germination of an early diverging land plant, the bryophyte *Physcomitrella patens*, in a lab-based assay. Low (<1µM) concentrations of AHLs promote spore germination whilst higher concentrations (5-10µM) inhibit spore germination. In general, AHLs with longer chain length (C8-C12) have a more potent effect than C4-C6 AHLs and side-group substitutions change the potency of germination-promoting activity with 3-O and 3-OH substitutions generally showing a slight reduction in potency.

The inhibitory effect of higher concentrations of AHLs is reminiscent of their effect in the green seaweed *Ulva* where >5µM AHLs can inhibit the early development and growth of new plants from zoospores ^69^. Higher concentrations (25-125µM) of long-chain AHLs can also inhibit *Ulva* spore swimming speed to promote settlement with 3-O substitutions showing the greatest inhibition ^71^. The effect of sub-micromolar concentrations AHLs was not investigated in these experiments.

In land plants, the effect of AHLs on germination of the desiccation-resistant dispersal units, namely spores (in Bryophytes, Lycophytes and ferns) or seeds (in Gymnosperms and Angiosperms) is not well studied. So far, a single study shows that priming of winter wheat (*Triticum aestivum* L.) seeds with C6-HSL (∼9ng AHL per seed) improves their germination and subsequent growth, development and biomass production ^76^. Thus, a potential role in germination control for AHLs is present across plant- and algal taxa, although whether this is as a result of convergent or divergent evolution is unknown. AHLs have a range of effects on post-germination development and growth in seed plants. For example, 1-10µM of 3O-C6-HSL and 3O-C8-HSL, and 10µM C4-HSL, C6-HSL, C8-HSL can increase *Arabidopsis* primary root elongation ^22,45,47,48,51^ while >10µM of C10-C14 AHLs inhibit primary root growth in *Arabidopsis* seedlings ^21,48^. Moreover C10 and C12 AHLs promote root branching and increases root hair formation at 12-96µM ^21^. Inhibitory effects of AHLs on the *Arabidopsis* root involve changes in cell division and differentiation ^21^. This biphasic pattern (growth stimulation of *Arabidopsis* primary root at low concentrations, growth inhibition at higher concentrations) is reminiscent of what we see with *Physcomitrella* spore germination (Figures 1-3). In general, longer-chain AHLs (C10, C12) have more potent effects, as we saw with *Physcomitrella* spore germination in this paper, although the concentrations required for an effect in *Arabidopsis* are higher (≥1µM) than in *Physcomitrella* (2nM-1µM). In barley, 10µM C6-HSL promotes seedling growth ^35^. Several studies hint at the molecular mechanisms underlying the effects of AHLs on *Arabidopsis* root growth. Transcriptome- and qRT-PCR approaches coupled with mutant studies implicate several transcription factors in the response, including *At*MYB44 ^51^, in addition to G-protein signalling ^47,48^ and calmodulin/calcium signalling ^45,46^. Interestingly, a role for changes in intracellular calcium signalling has also been implicated in *Ulva* spore settlement, which is also affected by AHLs ^77^.

There is considerable overlap between the bacteria we isolated from *Physcomitrella* sporophytes and the bacteria found in assocated with the peat moss *Sphagnum* ^56^ in which *Pseudomonas, Rahnella, Serratia, Stenotrophomonas* and *Microbacterium* are all present but Aeromonas and Acinetobacter were not detected. No *Bacillus* was detected in *Sphagnum*, although *Paenibacillus* (Firmicutes) was, along with additional Beta-proteobacteria, Bacteroidetes, and Actinobacteria ^56^.

The bacteria isolated from mosses are generally different from those isolated from *Ulva*: predominantly Alpha-proteobacteria and Bacteroidetes, although *Microbacterium* has been isolated from all three species ^78–80^. Most of the genera of Gamma-proteobacteria isolated (*Pseudomonas, Serratia, Acinetobacter, Aeromonas*) are AHL producers ^26,81–84^. However, it is important to note that only a small fraction (<1%) of all bacteria that exist in a particular environment can be grown in the lab on standard growth media ^85^ so there may be many other AHL-producing bacteria associated with *Physcomitrella* in the wild.

Single-species AHL analysis showed that, as expected, many of the *Pseudomonas* isolates, most of the *Serratia* isolates and one of the *Aeromonas* isolates produced AHLs (Figure 4). Moreover, an isolate of *Stenotrophomonas* from each of the Chew Valley and Stocks reservoir sites also produced AHLs: this genus has not previously been found to make AHLs as it normally makes DSF-type quorum sensing molecules ^86^ which also induce growth promoting traits on plants ^87^. Two *Pseudomonas fluorescences* isolates from the Stocks reservoir showed AHL production even though, to our knowledge, no stains from this species have been reported before to produce these QS molecules. Unexpectedly, none of the individual isolates from Lindley produced AHLs (Figure 4) despite several attempts, suggesting that they have either lost the ability to produce these molecules or, under the *in vitro* growth conditions used, they only make AHLs below the lower limit of detection for the LC-MS/MS system used.

In summary, we have characterised for the first time the effect of bacterial quorum sensing molecules, AHLs, on the developmen of a non-flowering land plant, the moss *Physcomitrella patens*. AHLs promote *Physcomitrella* spore germination at sub-micromolar concentrations, but inhibit germination at higher concentrations, in a biphasic pattern reminiscent of the AHL effect on root growth in flowering plants. We have shown that a range of bacteria, some of which produce AHLs, are associated with *Physcomitrella* sporophytes isolated from the wild. Future research could include a metagenomic analysis to identify all bacteria (including those that are uncultivatable) associated with *Physcomitrella*, work with mutant strains of bacteria deficient in AHL production, analysis of calcium signalling in moss spores upon AHL application, isolation of moss mutant strains that cannot respond to AHLs, or transcriptomic/proteomic analysis of moss spores treated with AHLs.

## Materials and Methods

### Moss spore germination assays

Germination assays were carried out as in ^88^. Briefly, spores from at least 3 age-matched sporophytes were used within each assay with three sporophytes’ worth of spores used for every 10 Petri dishes (9 cm diameter). Sporophytes were bleached in groups of two to three in 1 ml 25% Parozone™ (Jeyes Group, Thetford, UK) for 10 min and then washed three times in 1 ml sterile distilled water (10 min each) in a sterile flow cabinet. The sporophytes were then crushed in 100–200 µl of sterile water to release the spores. Spores were diluted down in sufficient sterile distilled water to allow plating of 500 µl of spore solution per Petri dish. Spores were plated on cellophane-overlaid BCD moss growth medium (1mM MgSO_4_, 1.84mM KH_2_PO_4_, 10mM KNO_3_, 45µM FeSO_4_.7H_2_O, plus 1:1000 Hoagland’s A-Z Trace Element Solution), supplemented with 5 mM CaCl_2_ and 5 mM ammonium tartrate. Cellophane discs (A.A. Packaging Ltd, Preston, UK) were autoclaved wet and individually between sheets of filter paper for 15 min at 121°C, before use. Each data point included data from more than one plate and a minimum of 500 spores.

### Isolation of wild *Physcomitrella patens*

*Physcomitrella patens* growing wild in the UK was isolated from 3 sites: Chew Valley Lake (Somerset; ST5814 6053), Stocks Reservoir (Yorkshire; SD742562) and Lindley (Yorkshire; 44/217414). A small area of moss containing ∼40 individual sporulating plants each harbouring a single sporophyte was collected, and kept moist during transport to the lab, where samples were refrigerated prior to sporophyte harvesting.

### Isolation and purification of bacteria

Initially, 12 sporophytes from each location were placed on individual Luria broth (LB)-agar plates and bacteria were allowed to grow out from the sporophyte for 2 days at 28°C in the dark (lower temperatures favoured growth of fungal contamination). The majority of sporophytes were associated with bacteria that could be grown on LB-agar, giving rise to bacterial consortia. These consortia were further purified by taking them through 3-4 rounds of streaking (giving rise to multiple single colonies) as appropriate, growing on LB-agar at 28°C overnight to obtain multiple pure bacterial isolates (identifiable by morphology and colour) for gDNA isolation, sequence identification and AHL detection. Stock plates for each strain were generated from a single colony and colonies from these plates were inoculated into liquid culture to make permanent glycerol stocks in 25% glycerol, 75% LB.

### Bacterial identification

Bacterial isolates were identified to genus-, or where possible species-level. Bacterial cultures were grown in LB from single colonies and genomic DNA was extracted using a Qiagen Blood and Tissue DNeasy kit (Qiagen, Hilden, Germany) following the manufacturer’s instructions. Partial 16S rDNA fragments (∼2kb) were amplified from 10-30ng of genomic DNA by PCR using the forward primer 27F (AGA GTT TGA TCC TGG CTC AG) and reverse primer 1522R (AAG GAG GTG ATC CAG CCG CA). PCR was carried out using Velocity proofreading DNA polymerase (Bioline) according to manufacturer’s instructions. The PCR cycling conditions were a denaturation of 94°C for 2 min followed by 30 cycles of 94°C for 30 sec, 58 °C for 30 sec and 72°C for 1 min, then a final extension of 72°C for 5 min. PCR products were purified using a GeneJET PCR purification kit (Thermo Fisher) and were sequenced using both the forward and reverse primers via capillary sequencing on an ABI3730 machine (Applied BioSystems). Raw sequence reads viewed in SnapGene version 1.4 and were trimmed and refined by eye from the peak trace where necessary. Where possible, forward and reverse sequences were aligned and combined to generate a single consensus sequence. Sequences were analysed by BLASTN ^89^ and the closest matches recorded.

### AHL analysis of bacteria

Bacterial cultures were grown in 5 ml of LB for 24 hr at 30°C with shaking at 200 rpm. For each sample, 1ml of filter sterilized supernatant was spiked with 5 µl of a 10µM solution of a deuterated AHL internal standard (d9-C5-AHL in MeOH). After solvent extraction (x3) with 0.5 ml aliquots of acidified ethyl acetate (0.1% (v/v) AcOH in EtOAc), combined extracts were dried under vacuum and stored at -20° prior to analysis. Dried samples were re-dissolved in 50µl of MeOH and 5.0µl of each sample injected for analysis.

For the analysis by LC-MS/MS, chromatography was achieved using a Shimadzu series 10AD LC system. The LC column, maintained at 40°C, was a Phenomenex Gemini C18 (3.0 µm, 100 x 3.0 mm). Mobile phases A and B were 0.1% (v/v) formic acid in water and methanol respectively. The flow rate throughout the chromatographic separation was 450µL/min. The binary gradient initially began at 10% B for 1.0 min, increased linearly to 50% B over 0.5 min, then to 99% B over 4.0 min. This composition remained for 1.5 min, decreased to 10% B over 0.1 min, and stayed at this composition for a 2.9 min period of re-equilibration.

For the MS detection of eluting AHLs, an Applied Biosystems Qtrap 4000 hybrid triple-quadrupole linear ion trap mass spectrometer equipped with an electrospray ionisation (ESI) interface was used. Analysis was conducted with the MS operating in positive electrospray (+ES) multiple reaction monitoring (MRM) mode, screening the LC eluent for specific unsubstituted, 3-O and 3-OH AHLs with even numbered acyl chain length from 4-14 carbons long, and the deuterated internal standard, comparing the retention time of detected analytes with authentic synthetic standards.

## Acknowledgments

We thank Jeff Duckett and Sharon Pilkington for wild *Physcomitrella patens* samples; Zaynub Chaudhry and Xulyu Cao for lab assistance. We thank the University of Birmingham Genomics Facility for their sequencing service. The work was funded by a UK-Natural Environment Research Council (NERC) PhD Studentship for EFV (NE/I528218/1), a grant from the Birmingham-Nottingham Strategic Collaboration fund (BNSCF002) to JC/MG and University of Birmingham funding for ALW, SN, KJ and WT to carry out Masters projects. This work was also supported by funding from the Biotechnology and Biological Sciences Research Council (BBSRC; Award Number BB/R012415/1). MC is partly funded by the National Biofilms Innovation Centre (NBIC) which is an Innovation and Knowledge Centre funded by the Biotechnology and Biological Sciences Research Council, Innovate UK and Hartree Centre.

## Author Contributions

EFV, MC, XL and JCC conceived and designed the study; EFV, ALW, SN, WT, KJ, NH, FG, XL and JCC performed experiments in the lab; EFV, ALW, SN, WT, KJ, NH and FG analysed data; EFV, SN, ALW, NH, FG, MC and JCC wrote sections of the manuscript. NH, MC and JCC revised and finalised the manuscript. All authors read and approved the submitted version.

## Additional information

### Accession Codes

All sequence data generated in this study has been deposited at GenBank and the accession numbers are given in Table 1.

### Data Availability Statement

The raw data supporting the conclusions of the germination assays and AHL quantification will be made available by the authors, without undue reservation, to any qualified researcher.

## References

1. Oldroyd, G. E. D. Speak, friend, and enter: Signalling systems that promote beneficial symbiotic associations in plants. Nat. Rev. Microbiol. 11, 252–263 (2013).

2. Mendes, R., Garbeva, P. & Raaijmakers, J. M. The rhizosphere microbiome: Significance of plant beneficial, plant pathogenic, and human pathogenic microorganisms. FEMS Microbiol. Rev. 37, 634–663 (2013).

3. Ortíz-Castro, R., Contreras-Cornejo, H. A., Macías-Rodríguez, L. & López-Bucio, J. The role of microbial signals in plant growth and development. Plant Signal. Behav. 4, 701–712 (2009).

4. Humphreys, C. P. et al. Mutualistic mycorrhiza-like symbiosis in the most ancient group of land plants. Nat. Commun. 1, 103 (2010).

5. Field, K. J. et al. First evidence of mutualism between ancient plant lineages (Haplomitriopsida liverworts) and Mucoromycotina fungi and its response to simulated Palaeozoic changes in atmospheric CO2. New Phytol. 205, 743–756 (2015).

6. Buhian, W. P. & Bensmihen, S. Mini-review: nod factor regulation of phytohormone signaling and homeostasis during rhizobia-legume symbiosis. Front. Plant Sci. 9, (2018).

7. Svistoonoff, S., Hocher, V. & Gherbi, H. Actinorhizal root nodule symbioses: What is signalling telling on the origins of nodulation? Curr. Opin. Plant Biol. 20, 11–18 (2014).

8. Hocher, V. et al. Signalling in actinorhizal root nodule symbioses. Antonie van Leeuwenhoek, Int. J. Gen. Mol. Microbiol. 112, 23–29 (2019).

9. Backer, R. et al. Plant growth-promoting rhizobacteria: Context, mechanisms of action, and roadmap to commercialization of biostimulants for sustainable agriculture. Front. Plant Sci. 9, 1–16 (2018).

10. Hartmann, A. & Schikora, A. Quorum Sensing of Bacteria and Trans-Kingdom Interactions of N-Acyl Homoserine Lactones with Eukaryotes. J. Chem. Ecol. 38, 704–713 (2012).

11. Schikora, A., Schenk, S. T. & Hartmann, A. Beneficial effects of bacteria-plant communication based on quorum sensing molecules of the N-acyl homoserine lactone group. Plant Mol. Biol. 90, 605–612 (2016).

12. Schenk, S. T. et al. N-acyl-homoserine lactone primes plants for cell wall reinforcement and induces resistance to bacterial pathogens via the salicylic acid/oxylipin pathway. Plant Cell 26, 2708–2723 (2014).

13. Cha, C., Gao, P., Chen, Y. C., Shaw, P. D. & Farrand, S. K. Production of acyl-homoserine lactone quorum-sensing signals by gram-negative plant-associated bacteria. Mol. Plant-Microbe Interact. 11, 1119–1129 (1998).

14. Pierson, L., Wood, D. & Beck von Bodman, S. Quorum sensing in plant-associated bacteria. in Cell-Cell Signalling (eds. Dunney, G. & Winans, S.) 101–116 (American Society for Microbiology Press, 1999).

15. Loh, J., Pierson, E. A., Pierson, L. S., Stacey, G. & Chatterjee, A. Quorum sensing in plant-associated bacteria. Curr. Opin. Plant Biol. 5, 285–90 (2002).

16. Whitehead, N. & Barnard, A. Quorum-sensing in Gram-negative bacteria. FEMS Microbiol. 25, 365–404 (2001).

17. Von Bodman, S. B., Bauer, W. D. & Coplin, D. L. Quorum sensing in plant-pathogenic bacteria. Annu Rev Phytopathol 41, 455–82 (2003).

18. Ramey, B. E., Koutsoudis, M., Bodman, S. B. Von & Fuqua, C. Biofilm formation in plant-microbe associations. Curr. Opin. Microbiol. 7, 602–9 (2004).

19. Whiteley, M., Diggle, S. P. & Greenberg, E. P. Progress in and promise of bacterial quorum sensing research. Nature 551, 313–20 (2017).

20. Mathesius, U. et al. Extensive and specific responses of a eukaryote to bacterial quorum-sensing signals. Proc. Natl. Acad. Sci. U. S. A. 100, 1444–1449 (2003).

21. Ortíz-Castro, R., Martínez-Trujillo, M. & López-Bucio, J. N-acyl-L-homoserine lactones: A class of bacterial quorum-sensing signals alter post-embryonic root development in Arabidopsis thaliana. Plant, Cell Environ. 31, 1497–1509 (2008).

22. Von Rad, U. et al. Response of Arabidopsis thaliana to N-hexanoyl-DL-homoserine-lactone, a bacterial quorum sensing molecule produced in the rhizosphere. Planta 229, 73–85 (2008).

23. Schikora, A. et al. N-acyl-homoserine lactone confers resistance toward biotrophic and hemibiotrophic pathogens via altered activation of AtMPK6. Plant Physiol. 157, 1407–18 (2011).

24. Bai, X., Todd, C. D., Desikan, R., Yang, Y. & Hu, X. N-3-oxo-decanoyl-l-homoserine-lactone activates auxin-induced adventitious root formation via hydrogen peroxide- and nitric oxide-dependent cyclic GMP signaling in mung bean. Plant Physiol. 158, 725–736 (2012).

25. Williams, P., Winzer, K., Chan, W. C. & Cámara, M. Look who’s talking: Communication and quorum sensing in the bacterial world. Philos. Trans. R. Soc. B Biol. Sci. 362, 1119–1134 (2007).

26. Williams, P. & Cámara, M. Quorum sensing and environmental adaptation in Pseudomonas aeruginosa: a tale of regulatory networks and multifunctional signal molecules. Curr. Opin. Microbiol. 12, 182–191 (2009).

27. González, J. F. & Venturi, V. A novel widespread interkingdom signaling circuit. Trends Plant Sci. 18, 167–174 (2013).

28. Scott, R. A. et al. Long- and short-chain plant-produced bacterial N-acyl-homoserine lactones become components of phyllosphere, rhizophere, and soil. Mol. Plant-Microbe Interact. 19, 227–39 (2006).

29. Churchill, M. E. A. & Chen, L. Structural basis of acyl-homoserine lactone-dependent signaling. Chem. Rev. 111, 68–85 (2011).

30. Qian, G., Xu, F., Venturi, V., Du, L. & Liu, F. Roles of a solo LuxR in the biological control agent Lysobacter enzymogenes strain OH11. Phytopathology 104, 224–31 (2014).

31. Hudaiberdiev, S. et al. Census of solo LuxR genes in prokaryotic genomes. Front. Cell. Infect. Microbiol. 5, 20 (2015).

32. Rowe, S. L., Norman, J. S. & Friesen, M. L. Coercion in the evolution of plant–microbe communication: A perspective. Mol. Plant-Microbe Interact. 31, 789–794 (2018).

33. Palmer, A. G., Senechal, A. C., Mukherjee, A., Ané, J. M. & Blackwell, H. E. Plant responses to bacterial N-acyl l-homoserine lactones are dependent on enzymatic degradation to l-homoserine. ACS Chem. Biol. 9, 1834–1845 (2014).

34. Calatrava-Morales, N., McIntosh, M. & Soto, M. J. Regulation mediated by N-acyl homoserine lactone quorum sensing signals in the rhizobium-legume symbiosis. Genes (Basel). 9, 263 (2018).

35. Klein, I., Von Rad, U. & Durner, J. Homoserine lactones: Do plants really listen to bacterial talk? Plant Signal. Behav. 4, 50–51 (2009).

36. Zarkani, A. A. et al. Homoserine lactones influence the reaction of plants to rhizobia. Int. J. Mol. Sci. 14, 17122–17146 (2013).

37. Sieper, T. et al. N-acyl-homoserine lactone uptake and systemic transport in barley rest upon active parts of the plant. New Phytol. 201, 545–555 (2014).

38. Gao, M., Teplitski, M., Robinson, J. B. & Bauer, W. D. Production of substances by Medicago truncatula that affect bacterial quorum sensing. Mol. Plant-Microbe Interact. 16, 827–834 (2003).

39. Bauer, W. D. & Mathesius, U. Plant responses to bacterial quorum sensing signals. Curr. Opin. Plant Biol. 7, 429–433 (2004).

40. Pérez-Montaño, F. et al. Rice and bean AHL-mimic quorum-sensing signals specifically interfere with the capacity to form biofilms by plant-associated bacteria. Res. Microbiol. 164, 749–760 (2013).

41. Corral-Lugo, A., Daddaoua, A., Ortega, A., Espinosa-Urgel, M. & Krell, T. Rosmarinic acid is a homoserine lactone mimic produced by plants that activates a bacterial quorum-sensing regulator. Sci. Signal. 9, ra1 (2016).

42. Nievas, F., Vilchez, L., Giordano, W. & Bogino, P. Arachis hypogaea L. produces mimic and inhibitory quorum sensing like molecules. Antonie van Leeuwenhoek, Int. J. Gen. Mol. Microbiol. 110, 891–902 (2017).

43. Manefield, M. et al. Halogenated furanones inhibit quorum sensing through accelerated LuxR turnover. Microbiology 148, 1119–27 (2002).

44. Koh, C. L. et al. Plant-derived natural products as sources of anti-quorum sensing compounds. Sensors (Switzerland) 13, 6217–6228 (2013).

45. Zhao, Q. et al. Involvement of calmodulin in regulation of primary root elongation by N-3-oxo-octanoyl homoserine lactone in Arabidopsis thaliana. Front. Plant Sci. 5, 807 (2015).

46. Song, S., Jia, Z., Xu, J., Zhang, Z. & Bian, Z. N-butyryl-homoserine lactone, a bacterial quorum-sensing signaling molecule, induces intracellular calcium elevation in Arabidopsis root cells. Biochem. Biophys. Res. Commun. 414, 355–360 (2011).

47. Jin, G. et al. Two G-protein-coupled-receptor candidates, Cand2 and Cand7, are involved in Arabidopsis root growth mediated by the bacterial quorum-sensing signals N-acyl-homoserine lactones. Biochem. Biophys. Res. Commun. 417, 991–995 (2012).

48. Liu, F., Bian, Z., Jia, Z., Zhao, Q. & Song, S. The GCR1 and GPA1 participate in promotion of Arabidopsis primary root elongation induced by N-Acyl-homoserine lactones, the bacterial quorum-sensing signals. Mol. Plant-Microbe Interact. 25, 677–683 (2012).

49. Miao, C., Liu, F., Zhao, Q., Jia, Z. & Song, S. A proteomic analysis of Arabidopsis thaliana seedling responses to 3-oxo-octanoyl-homoserine lactone, a bacterial quorum-sensing signal. Biochem. Biophys. Res. Commun. 427, 293–298 (2012).

50. Schuhegger, R. et al. Induction of systemic resistance in tomato by N-acyl-L-homoserine lactone-producing rhizosphere bacteria. Plant, Cell Environ. 29, 909–918 (2006).

51. Zhao, Q. et al. AtMYB44 positively regulates the enhanced elongation of primary roots Induced by N-3-Oxo-Hexanoyl-homoserine lactone in Arabidopsis thaliana. Mol. Plant-Microbe Interact. 29, 774–785 (2016).

52. Lugtenberg, B. J. J., Dekkers, L. & Bloemberg, G. V. Molecular determinants of rhizosphere colonization by Pseudomonas. Annu. Rev. Phytopathol. 39, 461–90 (2001).

53. Savka, M. A., Dessaux, Y., Oger, P. & Rossbach, S. Engineering bacterial competitiveness and persistence in the phytosphere. Mol. Plant-Microbe Interact. 15, 866–74 (2002).

54. Steidle, A. et al. Visualization of N-Acylhomoserine Lactone-Mediated Cell-Cell Communication between Bacteria Colonizing the Tomato Rhizosphere. Appl. Environ. Microbiol. 67, 5761–5770 (2001).

55. Joseph, C. M. & Phillips, D. A. Metabolites from soil bacteria affect plant water relations. Plant Physiol. Biochem. 41, 189–192 (2003).

56. Kostka, J. E. et al. The Sphagnum microbiome: New insights from an ancient plant lineage. New Phytol. 211, 57–64 (2016).

57. Glime, J. Nutrient Relations: CO2. in Bryophyte Ecology: Volume 1, Physiological Ecology (Ebook sponsored by Michigan Technological University and the International Association of Bryologists., 2017).

58. Glime, J. Household and Personal Uses. in Bryophyte Ecology: Volume 5, Uses (Ebook sponsored by Michigan Technological University and the International Association of Bryologists., 2017).

59. Glime, J. Medical Uses: Medical Conditions. in Bryophyte Ecology: Volume 5, Uses (Ebook sponsored by Michigan Technological University and the International Association of Bryologists., 2017).

60. Glime, J. Medical Uses: Biologically Active Substances. in Bryophyte Ecology: Volume 5, Uses (Ebook sponsored by Michigan Technological University and the International Association of Bryologists, 2017).

61. Bragina, A. et al. The Sphagnum microbiome supports bog ecosystem functioning under extreme conditions. Mol. Ecol. 23, 4498–4510 (2014).

62. Weston, D. J. et al. The Sphagnome Project: enabling ecological and evolutionary insights through a genus-level sequencing project. New Phytol. 217, 16–25 (2018).

63. Weston, D. J. et al. Sphagnum physiology in the context of changing climate: Emergent influences of genomics, modelling and host-microbiome interactions on understanding ecosystem function. Plant, Cell Environ. 38, 1737–1751 (2015).

64. Aschenbrenner, I. A., Cernava, T., Erlacher, A., Berg, G. & Grube, M. Differential sharing and distinct co-occurrence networks among spatially close bacterial microbiota of bark, mosses and lichens. Mol. Ecol. 26, 2826–2838 (2017).

65. Opelt, K. & Berg, G. Diversity and antagonistic potential of bacteria associated with bryophytes from nutrient-poor habitats of the baltic sea coast. Appl. Environ. Microbiol. 70, 6569–6579 (2004).

66. Tian, Y. & Li, Y. H. Comparative analysis of bacteria associated with different mosses by 16S rRNA and 16S rDNA sequencing. J. Basic Microbiol. 57, 57–67 (2017).

67. Hornschuh, M., Grotha, R. & Kutschera, U. Epiphytic bacteria associated with the bryophyte Funaria hygrometrica: Effects of Methylobacterium strains on protonema development. Plant Biol. 4, 682–687 (2002).

68. Kutschera, U. Plant-associated methylobacteria as co-evolved phytosymbionts: A hypothesis. Plant Signal. Behav. 2, 74–78 (2007).

69. Twigg, M. S., Tait, K., Williams, P., Atkinson, S. & Cámara, M. Interference with the germination and growth of Ulva zoospores by quorum-sensing molecules from Ulva-associated epiphytic bacteria. Environ. Microbiol. 16, 445–453 (2014).

70. Singh, R. P., Baghel, R. S., Reddy, C. R. K. & Jha, B. Effect of quorum sensing signals produced by seaweed-associated bacteria on carpospore liberation from Gracilaria dura. Front. Plant Sci. 6, 117 (2015).

71. Wheeler, G. L., Tait, K., Taylor, A., Brownlee, C. & Joint, I. Acyl-homoserine lactones modulate the settlement rate of zoospores of the marine alga Ulva intestinalis via a novel chemokinetic mechanism. Plant, Cell Environ. 29, 608–618 (2006).

72. Joint, I., Tait, K. & Wheeler, G. Cross-kingdom signalling: Exploitation of bacterial quorum sensing molecules by the green seaweed Ulva. Philos. Trans. R. Soc. B Biol. Sci. 362, 1223–1233 (2007).

73. Tait, K. et al. Disruption of quorum sensing in seawater abolishes attraction of zoospores of the green alga Ulva to bacterial biofilms. Environ. Microbiol. 7, 229–240 (2005).

74. Cove, D. The Moss Physcomitrella patens. Annu. Rev. Genet. 39, 339–358 (2005).

75. Prigge, M. J. & Bezanilla, M. Evolutionary crossroads in developmental biology: Physcomitrella patens. Development 137, 3535–3543 (2010).

76. Moshynets, O. V. et al. Priming winter wheat seeds with the bacterial quorum sensing signal N-hexanoyl-L-homoserine lactone (C6-HSL) shows potential to improve plant growth and seed yield. PLoS One 14, e0209460 (2019).

77. Thompson, S. E. M. et al. Membrane recycling and calcium dynamics during settlement and adhesion of zoospores of the green alga Ulva linza. Plant, Cell Environ. 30, 733–744 (2007).

78. Tait, K. et al. Turnover of quorum sensing signal molecules modulates cross-kingdom signalling. Environ. Microbiol. 11, 1792–802 (2009).

79. Burke, C., Thomas, T., Lewis, M., Steinberg, P. & Kjelleberg, S. Composition, uniqueness and variability of the epiphytic bacterial community of the green alga Ulva australis. ISME J. 5, 590–600 (2011).

80. Ghaderiardakani, F., Coates, J. C. & Wichard, T. Bacteria-induced morphogenesis of Ulva intestinalis and Ulva mutabilis (Chlorophyta): a contribution to the lottery theory. FEMS Microbiol. Ecol. 93, fix094 (2017).

81. Jung, B. K. et al. Genomic and phenotypic analyses of Serratia fonticola strain GS2: a rhizobacterium isolated from sesame rhizosphere that promotes plant growth and produces N-acyl homoserine lactone. J. Biotechnol. 241, 158–162 (2017).

82. Buddrus-Schiemann, K. et al. Analysis of N-acylhomoserine lactone dynamics in continuous cultures of Pseudomonas putida IsoF by use of ELISA and UHPLC/qTOF-MS-derived measurements and mathematical models. Anal. Bioanal. Chem. 406, 6373–6383 (2014).

83. Erdönmez, D., Rad, A. Y. & Aksöz, N. Quorum sensing molecules production by nosocomial and soil isolates Acinetobacter baumannii. Arch. Microbiol. 199, 1325–1334 (2017).

84. Swift, S. et al. Quorum sensing in Aeromonas hydrophila and Aeromonas salmonicida: Identification of the Luxri homologs AhyRi and AsaRi and their cognate N-acylhomoserine lactone signal molecules. J. Bacteriol. 179, 5271–81 (1997).

85. Yamamoto, H. Viable but nonculturable state as a general phenomenon of non-sporeforming bacteria, and its modeling. J. Infect. Chemother. 6, 112–114 (2000).

86. Huedo, P. et al. Two different rpf clusters distributed among a population of Stenotrophomonas maltophilia clinical strains display differential diffusible signal factor production and virulence regulation. J. Bacteriol. 196, 2431–2442 (2014).

87. Alavi, P. et al. The DSF Quorum Sensing System Controls the Positive Influence of Stenotrophomonas maltophilia on Plants. PLoS One 8, e67103 (2013).

88. Vesty, E. F. et al. The decision to germinate is regulated by divergent molecular networks in spores and seeds. New Phytol. 211, 952–966 (2016).

89. Johnson, M. et al. NCBI BLAST: a better web interface. Nucleic Acids Res. 36, W5–W9 (2008).

