## Supplemental Figure 1 for "Cross-kingdom signalling regulates spore germination in the moss *Physcomitrella patens*"

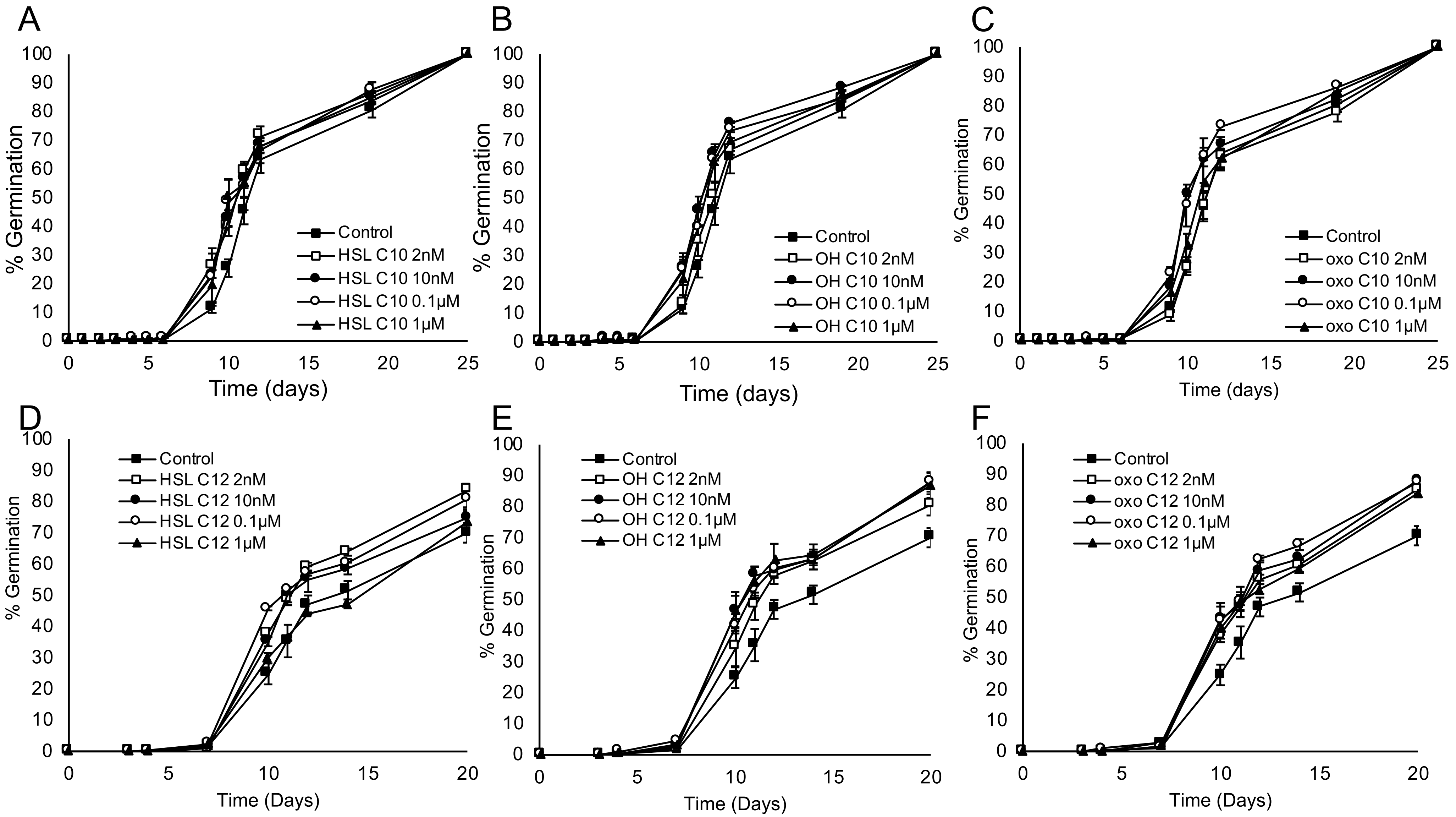


**Supplemental Figure 1. Effect of side group substitution on AHL activity towards *Physcomitrella* spore germination.**
A) Unsubstituted C10-HSL, B) 3-OH-C10-HSL, C) 3-O-C10-HSL, D) Unsubstituted C12-HSL, E) 3-OH-C12-HSL, F) 3-O-C12-HSL. These data are summarized in Figure 3B.
